# It is hard to be small: Inbreeding depression depends on the body size in a threatened songbird

**DOI:** 10.1101/2024.04.21.590470

**Authors:** Justyna Kubacka, Larissa S. Arantes, Magdalena Herdegen-Radwan, Tomasz S. Osiejuk, Sarah Sparmann, Camila J. Mazzoni

**Affiliations:** Museum and Institute of Zoology of the Polish Academy of Sciences, Twarda 51/55, 00-818 Warszawa, Poland; Berlin Center for Genomics in Biodiversity Research, Königin-Luise-Straße 6-8, 14195 Berlin, Germany; Leibniz Institute for Zoo and Wildlife Research, Alfred-Kowalke-Straße 17, 10315 Berlin, Germany; Department of Behavioural Ecology, Faculty of Biology, Adam Mickiewicz University, Uniwersytetu Poznańskiego 6, 61-614 Poznań, Poland; Leibniz Institute of Freshwater Ecology and Inland Fisheries, Müggelseedamm 310, 12587 Berlin Berlin, Germany

## Abstract

While inbreeding is known to affect individual fitness and thus population extinction risk, studies have under-represented non-model species of conservation concern, and rarely sought conditionality of inbreeding depression. Here, using SNPs identified with RAD-seq, we determined inbreeding depression in a threatened bird, the aquatic warbler *Acrocephalus paludicola*, and whether its magnitude depends on phenotypic and environmental factors. We found that the inbreeding coefficient (*F*) of adults with small tarsi was negatively associated with the seasonal breeding success (in males) and clutch size (in females), with the respective decrease in fitness in the most inbred relative to the least inbred individuals of ~89% and ~12%. In contrast, in adult males, for the average tarsus, wing and mass, support was low for *F* to be related to the long-term return rate to breeding grounds. For mean phenotypic covariates and male density, we also found low evidence that *F* is associated with the annual breeding success. Likewise, there was little support that mother *F* is related to egg hatch success and nestling survival, and – for average phenotypic traits, rainfall, temperature and nest density, and accounting for breeding peak – to clutch and fledged brood sizes. For nestlings, animal models showed that F is more negatively related to tarsus under higher temperatures and its effect varies by study year. However, for average brood size, temperature, rainfall and prey abundance, and when controlling for nestling sex, breeding peak and mother *F*, evidence for nestling F and tarsus association was weak. We conclude that (1) inbreeding depression on fitness components is stronger in smaller-bodied individuals, (2) considering interaction with phenotypic and environmental variables enables more accurate estimation of inbreeding depression, and (3) the inbreeding depression estimates will inform extinction risk analysis and conservation actions for the aquatic warbler.

## Introduction

Inbreeding results from mating between kin, non-random mating or division of a population into isolated groups (Keller & Waller 2002). It leads to increased genomic homozygosity and accumulation of harmful mutations, which are typically recessive and thus more often expressed in highly homozygous, inbred individuals. Consequently, more inbred individuals have compromised fitness. Inbreeding depression is found across plant and animal taxa (Ralls *et al*. 1988, Keller & Waller 2002) and is especially strong in wild populations (Crnokrak & Roff 1999). Inbreeding negatively affects key fitness components, such as sperm quality (Gage *et al*. 2006, Hinkson & Poo 2020), embryo development (Noordwijk & Scharloo 1981, Kruuk *et al*. 2002, Hemmings *et al*. 2012a), juvenile survival (Kruuk *et al*. 2002, Kennedy *et al*. 2014), annual reproductive success (Huisman *et al*. 2016, Niskanen *et al*. 2020), and lifetime reproductive success (Grueber *et al*. 2010, Huisman *et al*. 2016, Harrisson *et al*. 2019). Therefore, inbreeding depression decreases population viability and contributes to extinction risk (Westemeier *et al*. 1998, Brook *et al*. 2002, O’Grady *et al*. 2006, Nonaka *et al*. 2019), and evaluating its impact is of particular importance in populations of conservation concern.

Inbreeding depression is expected to be more pronounced in harsher environments or individuals of lower condition (Armbruster & Reed 2005). For example, low food availability and high number of competitors enhance the inbreeding depression on survival in Darwin’s finches (*Geospiza sp*.) (Keller *et al*. 2002). A long-term study on the great tit (*Parus major*) found that inbreeding by environment interactions, while generally weak, are stronger in adverse conditions more closely related to fitness (Szulkin & Sheldon 2007). Laboratory experiments on insects showed that inbreeding effects on offspring survival increase with mother age (Fox & Reed 2010) and environmental toxicity (Nowak *et al*. 2007). However, evaluation of the conditionality of inbreeding depression in wild populations has not been frequent, while determining its magnitude across individual traits and environmental conditions would yield more accurate inbreeding depression estimates, especially in threatened populations.

Here, we determine the strength of inbreeding depression on fitness-related traits in an IUCN-threatened songbird, the aquatic warbler *Acrocephalus paludicola*, a habitat specialist breeding in fen mires and a long-distance migrant (Le Nevé *et al*. 2018, Tanneberger *et al*. 2018b). Following the gradual disappearance of fens, which accelerated especially over the past 100-200 years due to peat extraction, changes in hydrological regime, drainage, eutrophication and succession, the extent of occurrence of the species shrank considerably (Tanneberger *et al*. 2018a). This led to a steep decline in the population size, amounting to 95% only between 1950-80, and its extinction from the western Europe by the end of the 20th century (Briedis & Keišs 2016, Flade *et al*. 2018). Today, the breeding range of the aquatic warbler is constrained to the east-central Europe, which holds c. 98% of the global population estimated at 12200 singing males (Flade *et al*. 2018). A recent study (Kubacka *et al*. 2024) demonstrated that genetic diversity in the species is low and comparable to that of other threatened bird species (Evans & Sheldon 2008, Vitorino *et al*. 2019). As it is habitat specialists that are more strongly affected by loss of genetic diversity (Matthews *et al*. 2014, Pflüger *et al*. 2019), it is crucial to determine the magnitude of inbreeding depression in the aquatic warbler. In addition, alongside the depleted genetic diversity, the high asymmetry of the male reproductive success (Dyrcz *et al*. 2002, 2005, Kubacka *et al*. 2025) and the promiscuous mating system (Leisler & Schulze-Hagen 2011) are expected to increase variation in the inbreeding rates, which facilitates detection of inbreeding depression (Balloux *et al*. 2004), thus making the species a good study model. Finally, the aquatic warbler faces a broad range of environmental conditions, as the breeding season strides the cool spring and the hot summer, water levels on the fen are changeable, and nests are built on the ground and thus exposed to environmental factors, all of which enable studying the conditionality of inbreeding depression.

Specifically, we aimed at 1) assessing the relationship of adult inbreeding rates with the reproductive success and survival, and of nestling inbreeding rates with the body size; 2) evaluating whether inbreeding depression is conditioned by phenotypic variables corresponding to individual quality and by environmental variables that are expected to affect adult survival and fecundity, and offspring body size; and 3) quantifying the inbreeding load and change in fitness associated with inbreeding.

## Methods

### Study area and sampling

The study was conducted between May-August of 2017-2023, in the Biebrza Valley (Poland), which holds about 25% of the global breeding population of the aquatic warbler. The breeding habitat is permanently water-logged open fen mire and to a less extent wet meadows, all dominated by sedges *Carex* spp. Three study areas were used, referred to as Ławki (N 53<u>º</u>17⍰11.4⍰⍰, E 22<u>º</u>33⍰49.2⍰⍰), Szorce (N 53<u>º</u>17⍰34.8⍰⍰, E 22<u>º</u>37⍰15.9⍰⍰) and Mścichy (53<u>º</u>25⍰41.7⍰⍰ E 22<u>º</u>30⍰17.0⍰⍰). Each study area consisted of two 10-20-hectare (ha) study plots, totalling 70 ha.

Between May-July of 2017-2019, adult individuals (past their 1^st^ calendar year) were caught with mist-nets and marked with unique metal and colour rings. They were blood-sampled through puncture of the brachial vein with a sterile needle and collecting approx. 10-120 ul of blood with a capillary. Each blood sample was immediately placed in a vial with an o-ring seal, containing 2 ml of 96% EtOH, and shaken to avoid formation of large clots. We measured tarsus length (from the notch on the metatarsus to the top of the bone above the folded toes, with a calliper to the nearest 0.1 mm), wing length (with a ruler to the nearest mm) and body mass (with an electronic Kern CM 150-1N balance to the nearest 0.1 g).

Nests were located by search and observation of alarming females, every 2-7 days in each plot from late May to early June, and from late June until the end of July. We distinguished two breeding peaks, which correspond to the first and second breeding attempts that are typically separated by about 10 days when no nests are initiated (Kubacka *et al*. 2014). Nests were monitored every 1-5 days. In nests found at the egg stage or on day 1 post-hatch (day 0 being the last day when most offspring in the nest were still eggs), we determined the clutch size, hatched brood size and nestling survival (number of chicks surviving from hatching to fledging). After day 1 the risk is higher that a nestling dies and is removed by the female, which biases estimation of the above traits. In all the nests, we recorded the fledged brood size as the number of nestlings seen on the last check when the brood was still in the nest, as long as on the following check the nest did not bear any traces of predation and was recorded empty not earlier than on day 12 post-hatch (i.e. 3 days before the expected fledging).

In 2017-2018, the tarsus and body mass of chicks were measured, as above, on days 2, 5 and 9 post-hatch, which fall at the beginning, in one third and in two thirds of the nest stage, respectively. Only nestlings from nests found at the egg stage or day 1-2 post-hatch were included in the analysis. We used the nestling tarsus length as it is one of the proxies for structural body size in birds (Rising & Somers 1989, Senar & Pascual 1997) and predicts survival to reproductive maturity (Gebhardt-Henrich & Richner 1998). Chicks were blood-sampled (approx. 10-80 μl) on day 9 post-hatch, as above.

Between May-July of 2018-2023, we searched the study plots and their surroundings up to 150 m off plot border for colour-ringed adults with the search effort of 0.2-3.9 person-hours per ha. The colour-ring code was read with an 80-mm-lens and 20-60 magnification scope. The position of each resighted individual was stored in a GPS receiver. We measured the long-term return rate by the number of years during which a given adult individual was resighted or recaught in the study plots, or in other breeding areas – as determined from the Polish Bird Ringing Database, over 4 years following the first observation as adult.

We obtained the daily sum of rainfall in mm and average daily temperature from the Polish Institute of Meteorology and Water Management (IMGW). To assess food abundance, in 2017-2018 we sampled arthropod prey within c. 30 m from the nest by sweep-netting (1 sweep per 1 second) on 3 radially oriented transects between 3 m of the nest and 50 steps away from the nest. Collected arthropods were placed in a plastic bag, frozen upon arrival from the field, determined to the order and counted. We then quantified the amount of preferred prey as the total number of insects from the taxonomic groups known to be selectively brought by females to nestlings: *Odonata, Aranea, Lepidoptera* and *Orthoptera* (Schulze-Hagen *et al*. 1989). We estimated the male density with QGIS 3.10.7 (QGIS Development Team 2020) as the average number of ringed males observed within 100 m of each ringed male’s position, and the nest density as the number of nests found per 10 ha of study plot. Following our previous study (Kubacka *et al*. 2025), based on all the captures and resights of the males, we determined the number of encounters of a male per plot in each breeding season, to account for excess zeros in the male breeding success (see *Statistical analysis*). We suspected that the low father-to-offspring assignment rate (see *Results*) could be caused mainly by late arrivals of some males to the study areas. Individuals with a low number of encounters were first captured during the second breeding attempt, or were rarely seen in the study plots.

### DNA extraction and molecular sex identification

DNA was isolated with the Xpure^™^ Blood Mini kit (A&A Biotechnology, Poland). 200-300 μl of blood suspended in 96% EtOH were used per individual and the proteinase K digestion time was 2-3 hours. The remaining steps were performed according to the manufacturer’s protocol. We analysed DNA of 175 adult males (1 was ringed as a chick), 320 chicks (2 were resighted as adult males) from 79 nests and 105 adult females. DNA concentration was measured with a Picogreen (Invitrogen) or Qubit (Thermo Fisher Scientific) instrument.

Sex of nestlings was determined by PCR amplification of sex-specific sequences in the CHD-W (c. 370 bp, females only) and CHD-Z genes (c. 330 bp, both sexes) (Griffiths *et al*. 1998) with a PCR kit (Eurx, Poland). Electrophoresis of PCR products ran for 90 mins at 100 V in a 3% agarose gel stained with Simply Safe (Eurx, Poland) and the gel was photographed under UV light.

### Molecular marker identification and genotyping

To determine inbreeding rates and assign paternity, we used single nucleotide polymorphisms (SNPs), determined with 3-enzyme restriction site-associated sequencing (3RAD) (Bayona-Vásquez *et al*. 2019, Glenn *et al*. 2019). Inbreeding estimated with genomic markers better predicts the identity by descent and decrease in fitness, compared to inbreeding rates inferred from the pedigree (Hemmings *et al*. 2012b, Kardos *et al*. 2015, Huisman *et al*. 2016). First, we constructed a reference genomic library, i.e. a catalogue of reliable loci excluding potential paralogs and artefactual alleles, according to the Reduced-Representation Single-Copy Orthologs (R2SCOs) method (Driller *et al*. 2021), with modifications. We sequenced 2×300 bp reads for three females to reconstruct the entire locus by merging overlapping reads, using samples with high DNA concentration (75.9, 53.7 and 76.6 ng/µl), each representing a different study location. After normalising the samples to 20 ng/µl, we split each into 5 digestion reactions (200 ng each), totalling 1 µg digested DNA. The enzymes Xbal, EcoRI-HF and Nhel were used for the digestion, which ran at 37°C for 2 hours. Each digestion reaction was ligated with a unique Xbal and EcoRI adapter pair (i.e. double-indexed) and the 5 reactions were pooled and purified (0.8x PCRclean magnetic beads - GCbiotech). We performed a PCR with P5 and P7 customised ligation check primers, to ensure that the digestion was successful. Each pooled sample was size-selected (350-580 bp) on a BluePippin machine (Sage Science). Each size selection product was then split into 4 reactions and in a single-cycle PCR we added an iTru5-8N index (with 8 random nucleotides, to mark unique DNA template molecules and remove PCR duplicates during the bioinformatic analysis) (Hoffberg *et al*. 2016). After purification (0.8x beads) an additional PCR step (8 cycles) completed the Illumina adapters with a unique iTru7 P7 primer per reaction and a P5 primer (which has a flow cell binding site sequence) common to all the reactions. The 4 PCR replicates of each sample were then pooled, purified (0.8x beads) and quantified (Qubit – Thermo Fisher Scientific). Finally, the pools were sequenced with the Illumina MiSeq V3 at 600 cycles, yielding a total of 3.6 M reads per sample.

Next, we constructed population libraries (Supplementary Online Resource 1), consisting of the remaining 597 samples. DNA concentration was normalised to 20 ng/ul or less (at least 8.14 ng/µl). We followed the protocol described above (Bayona-Vásquez *et al*. 2019), with modifications, and produced 9 population libraries, which were processed in 3 batches. The libraries were sequenced with 2×150 bp on Illumina instruments; 15% of PhiX was used to increase sequence diversity. The number of sequenced reads obtained from each library is presented in Table S1a.

### Bioinformatic analysis

We first constructed a reference locus catalogue using the R2SCOs method (Driller *et al*. 2021), which allowed us to select a set of high-quality single-copy orthologous loci to be used as reference for the population analysis. The R2SCOs pipeline included de-replication of the pre-processed and size-selected sequences (range 250-370 bp, for which we obtained comparable and high coverage), setting the minimum coverage per unique sequence to define a putative allele equal to 3; and clustering and definition of putative loci using the intra-specific identity threshold of 90%. As a next step, the pipeline applies a set of filters to remove loci that are prone to generate spurious results due to issues of either biological or technical background. Aiming to avoid sex-biased genotypes that might interfere with the population analyses (Faux *et al*. 2020), we removed all the R2SCOs loci that mapped to the sex chromosomes of the Eurasian blackcap (*Sylvia atricapilla*) genome (GenBank accession number GCA_009819655.1). The final reference contained 10997 loci.

The raw data from the population libraries was mapped against the PhiX genome to filter any remaining read from the Illumina library control. Next, we performed adapter trimming using the Cutadapt software (Martin 2011). Then, reads were attributed to individuals based on the in-line barcodes using FLEXBAR (Dodt *et al*. 2012). In part of the libraries (Supplementary Online Resource 1), PCR duplicates were filtered out using the Python script Filter_PCR_duplicates.py (https://github.com/BeGenDiv/Arantes_et_al_2020). The 4 replicates were then concatenated. The forward and reverse reads were merged with overlapping regions of at least 30 bp and maximum length of 249 using the software PEAR (Zhang *et al*. 2014). All the merged reads (<249 bp) were considered short fragments that were out of the target range, and we continued analysing the unassembled read pairs, which were filtered for minimum quality (Q>30) and trimmed to 142 bp with a minimum length of 130 bp using Trimmomatic (Bolger *et al*. 2014). We checked the presence of the XbaI and EcoRI restriction sites in the paired-read ends using the script checkRestrictionSites.py (https://github.com/BeGenDiv/Arantes_et_al_2020) and filtering out any fragment resulting from star activity or NheI digestion. Finally, reads containing the internal restriction site of XbaI, EcoRI or NheI were removed using the Filter_Reads.py script (https://github.com/BeGenDiv/Arantes_et_al_2020). The filtered reads were mapped against the reference locus catalog constructed with the R2SCOs method, using Bowtie2 (Langmead *et al*. 2009) with default parameters and the flags ‘-no-mixed’ and ‘-no-discordant’. In order to select a common range of high-coverage loci for all individuals belonging to different libraries, we checked the length distribution of the loci based on the .sam files, and extracted the range 250-310 bp with *samtools* (Danecek *et al*. 2021) and a bash *awk* command.

We analysed the mapped reads with Stacks (Catchen *et al*. 2011, 2013), using the reference-based pipeline. Genotypes were called and filtered with ref_*map*.*pl*, which uses the *gstacks* and *populations* modules. In *gstacks*, the *var_alpha* (alpha threshold for discovering SNPs) was set to 0.01, and all the samples were assumed to belong to one population, because of the extensive gene flow known in the aquatic warbler (Kubacka *et al*. 2024). Otherwise, default parameters were used. In populations, we used: the minimum percentage of individuals in a population required to process a locus r= 0.8, to avoid high missingness; the minimum number of populations a locus must be present in to be processed p= 1; and the minimum minor allele frequency required to process a nucleotide site at a locus min-maf= 0.005, which ensured that an allele is present in at least three samples to be processed (Rochette & Catchen 2017). To obtain independent SNPs, we wrote only the first SNP in a locus. The remaining parameters were retained at their defaults. After calling genotypes, we generated mean depth per individual with Vcftools (Danecek *et al*. 2011), to inspect coverage distribution. We removed 1 individual with mean depth <10 and repeated the SNP-calling step. We obtained 6267 loci, of which 4179 variant sites (SNPs) were retained after the *populations* filter, with mean ±SE observed heterozygosity in variable positions of 0.141 ±0.002. We checked whether individuals with the lowest coverage (mean depth of ≥10x and <30x, N= 46) affect loci-calling, by running *ref_map*.*pl* without them. The number of SNPs called without these individuals was 4171 and the mean ±SE observed heterozygosity was 0.143 ±0.002; we therefore retained them for further analysis.

We then used Vcftools for further SNP filtering (Table S1b). We removed genotypes with mean depth per SNP <10x and loci with >20% missing data. Next, we filtered out sites (N= 6) of mean depth >115, which formed the tail of the depth distribution, to remove potential collapsed paralogs. In the R environment (R Core Team 2024), with the *dartR* package (Gruber *et al*. 2018) we checked whether the loci are in the Hardy-Weinberg equilibrium (HWE), applying the false discovery rate correction and p-value of 0.05. Most loci not in HWE were heterozygote-deficient and we found a positive correlation between the locus Fis and missingness (r= 0.38), which decreased to r= 0.13 after the HWE-violating loci were removed. As this is indicative of null alleles, which artefactually lower heterozygosity (Waples 2015, De Meeûs 2018), we proceeded without the HWE-violating SNPs. Finally, using VCFtools, we calculated loci-pairwise *r*^2^ to inspect distribution of linkage disequilibrium. The mean ±SD of *r*^2^ was 0.003±0.012. We excluded one randomly selected SNP from each pair for which *r*^2^ was >0.3, to remove the tail of *r*^2^ distribution (Fig. S1). The final dataset consisted of 2948 SNPs (Table S1b). The mean ±SD read depth per locus and individual are provided in Fig. S2.

### Paternity assignment

We selected a set of informative SNPs to maximise the statistical power of parentage assignment (Andrews *et al*. 2018, Thrasher *et al*. 2018). With Vcftools (Danecek *et al*. 2011), we filtered out SNPs with minor allele frequency <0.3, as rare SNP variants are not informative in parentage assignment (Huisman 2017), and missingness >0.1. The number of SNPs after filtering was 333 (Table S1b). Typically, 100-200 SNPs suffice for parentage assignment (Huisman 2017, Flanagan & Jones 2019). For 1 individual, which was RAD-sequenced twice, we removed the genotype with fewer loci typed.

The resulting genotypes dataset consisted of 320 offspring (316 with known mother) and 173 candidate fathers, with 1 mother typed in 88 SNPs and all the other individuals typed in >99 SNPs. Paternity was then assigned with Cervus (Kalinowski *et al*. 2007), running the allele frequency analysis, parentage analysis simulation and paternity analysis given known mother. The allele frequency analysis showed the mean proportion of SNPs typed of 0.954. In the simulation, we assumed: the allele frequencies generated in the previous step; 100000 offspring; 440 as the average number of candidate fathers per offspring (based on the number of males assessed within the study plots and 150 m around them, and summed over all the three study locations); 0.4 as the proportion of candidate fathers sampled; 0.954 and 0.01 as the proportion of loci typed and mistyped, respectively; 0.01 as the error rate in likelihood calculations; 88 as the minimum number of typed loci; Delta as the statistic to determine confidence; and 99% and 95% as the strict and relaxed confidence level, respectively. In the paternity analysis, we used the allele frequency and simulation results generated in the previous steps. The offspring file contained candidate fathers, and males known to not be sexually mature in the year when a given chick was born were excluded. We ran the simulation and paternity analyses also for 95% and 80% strict and relaxed confidence levels, respectively. In addition, we established paternity with Colony v. 2.0.7.1 (Jones & Wang 2010), providing information on known maternal sibships and excluded paternity, assuming polygamy of both sexes, no inbreeding and default parameters otherwise. We quantified the male seasonal breeding success as the number of offspring produced in the given year, as detected by paternity assignment to the blood-sampled chicks.

### Inbreeding rate and inbreeding load

To assess inbreeding, we quantified the inbreeding coefficient *F* with a method of moments (Wang 2014) using the *–het* option in VCFtools (Danecek *et al*. 2011). To explore how well *F* corresponded to inbreeding, with the R package *inbreedR* (Stoffel *et al*. 2016), we calculated *g*_2_, an estimate of identity disequilibrium which quantifies the covariance of heterozygosity at the studied loci standardised by their average heterozygosity; hence, it corresponds to inbreeding variance in the population (David *et al*. 2007). A correlation between a trait and heterozygosity will not arise when *g*_2_= 0 (Szulkin *et al*. 2010). To avoid pseudoreplication, we used a dataset filtered to include all the adult individuals and 1 nestling per nest. We used 1000 permutations over SNPs to evaluate whether *g*_2_> 0 and 1000 bootstraps over individuals to determine its 95% confidence interval (CI).

To quantify the strength of inbreeding depression, we used the inbreeding load measured with lethal equivalents (*B*) and percent change in fitness due to inbreeding (*δ*) (Morton & Crow 1956, Keller & Waller 2002). Assuming that mutations at different loci have independent effects on fitness, the logarithm of a fitness-related trait is predicted to show a linear negative relationship with the inbreeding coefficient. The inbreeding load is quantified as the negative of the slope describing this relationship. To compare inbreeding load between populations and studies, the concept of lethal equivalents was coined. The effect of a mutation on fitness can be lethal or have partial detrimental effects. One lethal equivalent is the average number of alleles in a population which cause death of the homozygous individual. The inbreeding load equals the average number of lethal equivalents per haploid gamete found in the studied population. We calculated *B* using Equation II in Box 3 from Keller & Waller (2002), predictions weighted-averaged across all the models and categorical levels (see *Statistical analysis*), and fitness values for the maximum and minimum inbreeding rates in our dataset. For the survival traits, we used Poisson models with the logarithm link function, which yield unbiased estimates of inbreeding load for binomial traits (Nietlisbach *et al*. 2019). The *δ* refers to the change in a fitness-related trait in inbred individuals compared to outbred individuals (Keller & Waller 2002). We calculated *δ* for the most inbred compared to the least inbred individuals in the dataset (Harrisson *et al*. 2019, Sin *et al*. 2021).

### Statistical analysis

We ran the analysis in the R environment 4.4.2 (R Core Team 2024) and applied the information-theoretic approach, in which an a priori set of biologically plausible candidate models is built and each model in the set obtains a relative rank indicating how well it explains the data (Burnham & Anderson 2002). To rank models within a candidate set, we used the Akaike information criterion corrected for small sample size (AICc) and – in animal models (see below) – the deviance information criterion (DIC). In addition, we generated the following quantitative measures of relative support for each model: model likelihood (relative likelihood of the model in the candidate set, given the data), Akaike/DIC weight (ωAICc/ωDIC; probability that the given model is the best approximating model in the candidate set), evidence ratio (showing the probability of the best model relative to other models in the set) and cumulative AICc/DIC (sum of AICc/DIC weights of a given model and all the higher-ranking models) (Symonds & Moussalli 2011).

Each candidate set included a null model assuming that the response variable is constant, or that it varies with a variable strongly affecting the response that was common to all the models. All the models in the set except the null model included *F*. For selected candidate sets, we restricted the number of variables in a model to allow 10-15 data points per each non-intercept estimate and reduce the risk of over-parametrisation; for models using the binomial distribution, we ensured that there are 10-15 data points of the less probable event per estimate (Harrell 2015). To account for model uncertainty, the estimates of regression coefficients and their confidence intervals were weighted-averaged (by the ωAICc/ωDIC) across all the models in a set that contained the given term (i.e. natural model-averaging), as we expected small effects (Grueber *et al*. 2011, Symonds & Moussalli 2011). Model selection and averaging were carried out with package *AICmodavg* (Mazerolle 2023). All the input numerical explanatory variables were standardised to the mean of zero and SD of 1 (i.e. z-score standardisation). The weather variables, nest density and prey abundance were z-score standardised within the year and breeding peak; in the chick tarsus analysis, the weather variables were also standardised within the day. Sum contrasts were used (i.e. with the intercept being the corrected mean), to enable comparison of effects of categorical and numerical variables. For visualising interactions, predictions were calculated using packages *AICmodavg* (Mazerolle 2023) and *MuMIn* (Bartoń 2019) for the model containing the term, on the link scale and back-transformed. We visualised results with package *ggplot2* (Wickham 2016) and custom code.

We evaluated the relationship between *F* as an explanatory variable and the fitness-related (male long-term return rate, male breeding success, clutch size, hatch success, nestling survival and fledged brood size) and condition-related traits (chick tarsus on days 2, 5 and 9 post-hatch) as response. We also determined whether the *F* effect is conditioned by phenotypic (tarsus, wing and body mass in adults; sex in offspring) or environmental variables (male density, nest density, rainfall, temperature, breeding peak, prey abundance and brood size). Candidate sets were built as follows:

1. *Male inbreeding and long-term return rate*. We did not consider the female return rate due to only 8 recoveries, which did not ensure a reliable regression analysis (Harrell 2015). The male 4-year-return rate showed a right-skewed distribution. In order to select an appropriate count model, following the recommendations by Zuur & Ieno (2021), we ran an intercept-only GLM Poisson (*P*) model with log-link with the return rate as response, and calculated its overdispersion and zero-inflation using package *DHARMa* (Hartig 2022). The *P* model showed overdispersion (dispersion statistic 1.44, *P*<0.001) but no zero-inflation (ratio of observed to simulated zeros 1.09, *P*= 0.194). We then ran an intercept-only negative-binomial (NB) model using package *MASS* (Venables & Ripley 2002), which showed no overdispersion (1.00, *P*=0.997) and no zero-inflation (1.00, *P*= 1.000). Therefore, we continued with the NB distribution to construct a candidate model set. We included the following terms: body mass, tarsus and wing length, ringing area and year, their interaction with *F, F* squared, body mass squared, tarsus squared, and wing squared (Table S2a).
2. *Male inbreeding and seasonal breeding success*. The male breeding success had a large number of zeros and, corresponding to previous observations (Dyrcz *et al*. 2002, 2005, Kubacka *et al*. 2025), was right-skewed. We proceeded as in point (1) and constructed a null GLM *P* log-link model with the number of encounters and ringing year as explanatory variables. The *P* model was overdispersed (dispersion statistic 4.31, *P*<0.001) and zero-inflated (ratio of observed to simulated zeros 1.70, *P*<0.001). We then fitted a *NB* model with the same terms, which was not overdispersed (0.99, *P*= 0.965) and not zero-inflated (1.02, *P*= 0.786). Hence, we proceeded with the *NB* distribution. While 24/145 males bred both in 2017 and 2018, we treated these breeding events as independent after ascertaining that the residuals within these males were uncorrelated in a null GLM *NB* model (Zuur *et al*. 2009, 2013). The covariates included were body mass, tarsus and wing length, local male density, study area and ringing year, their interactions with *F*, and *F*, mass, wing and tarsus square effects. All the models included the number of encounters and ringing year (Table S2b).
3. *Female inbreeding and clutch size*. This candidate set was constructed using linear models. As covariates, we included the mother body mass, tarsus and wing length, and their quadratic effects, breeding peak (present in each model as it strongly predicts the clutch size), sum of rainfall during and mean daily temperature averaged for 15 days prior to start of incubation, study area and year, and the interactions of all the covariates with mother *F*. We restricted the number of non-intercept estimates in a model to 6 (Table S4a).
4. *Female inbreeding and egg hatching success*. The hatching success was coded as 1 for eggs that hatched and 0 otherwise, with the egg being the subject. The candidate set was built using generalized linear mixed models (GLMMs) with package *lme4* (Bates *et al*. 2015) and the binomial distribution with logit link. Nest ID was entered as a random factor. As covariates, we included the breeding peak, sum of rainfall and mean average daily temperature during incubation, clutch size, study area and year. Due to the low number of unhatched eggs (*N*= 23), we limited the number of non-intercept estimates in a model to 2 and did not consider interactions (Table S4b).
5. *Female inbreeding and nestling survival*. The nestling survival was coded as 1 for nestlings that survived until fledging and 0 otherwise, with the nestling being the subject. The candidate set was constructed with GLMMs, as above. As covariates, we included the mother body mass, wing and tarsus length, breeding peak, sum of rainfall and mean average daily temperature from hatching to fledging, hatched brood size, study area and year. Due to the low number of nestling deaths (*N*= 22), we limited the number of non-intercept estimates in a model to 2 and did not consider interactions (Table S4c).
6. *Female inbreeding and fledged brood size*. We considered only successful nests (i.e. those that fledged at least 1 chick), as nests fail completely for reasons that are mostly independent of the female (e.g. predation). The candidate set was built using linear models. As covariates, we included the mother body mass, wing and tarsus length and their quadratic effects, breeding peak, sum of rainfall and mean average daily temperature from hatching to fledging, study area and year. We also considered interactions of the covariates with *F*. The number of non-intercept estimates in a model was <8 (Table S4d).
7. *Nestling inbreeding and tarsus length on days 2, 5 and 9 post-hatch*. In natural populations, phenotype-associated inbreeding can occur, e.g. if the wing length of a parent is correlated with its relatedness to the mate. This creates a bias on the inbreeding depression especially in morphological traits, as part of variation in the trait results from genetic covariance with the relatedness between parents. Therefore, as recommended by Becker *et al*. (2016), we applied the animal model. We used package *MCMCglmm* (Hadfield 2010), with the random effects being the breeding value (‘animal’), chick ID, nest ID and father ID (from the Colony results). We calculated the genomic relatedness matrix using package *AGHmatrix* (Amadeu *et al*. 2023) and the Van Raden matrix. We applied an uninformative prior and set the number of MCMC iterations to 1.2 million, burnin to 10000 and thinning to 100. The candidate set considered the following covariates: measurement day (present in all models), chick sex, brood size, breeding peak, sum of rainfall during the previous 14 days, mean average daily temperature over the previous 4 days, prey abundance, area and year, and their interaction with chick *F*; and mother *F*, to account for an effect of maternal inbreeding on brood provisioning and parent-offspring correlation in *F* (Nietlisbach *et al*. 2016) (Table S6).

## Results

### Parentage assignment and inbreeding statistics

Of the 320 assignments, in further analysis we used 223 that were of ≥99% confidence in both the father-offspring pair and trio (i.e. father-mother-offspring), and 2 that were of ≥99% confidence in father-offspring pair, but their mother was not sampled and trio confidence could not be determined. We excluded 1 assignment for which the trio confidence was 99%, but the pair confidence was <95%. Of the 173 candidate fathers, 58 were assigned paternity with confidence as above. The range and median number of mismatches were: 0-3 and 0 for mother-chick; 0-5 and 1 for the 225 trios with the most likely father; and 0-49 and 39 for the 225 trios with the second most likely father. We obtained same assignments with the strict and relaxed confidence of 80% and 95%, respectively. All the Colony paternity assignments matched the 225 Cervus assignments and had the probability of 1.

The mean ±SD *F* was 0.005 ±0.043, median 0.002 and range –0.077 to 0.138 for adult females; and mean ±SD 0.028 ±0.048, median 0.030 and range –0.101 to 0.168 for adult males (Fig. S3). Two nestlings with unusually low inbreeding coefficients were removed from the tarsus analysis as outliers. The mean ±SD offspring *F*, excluding these outliers, was 0.007 ±0.046, median 0.006 and range –0.096 to 0.175. The *g*2 was estimated at 0.0009 (95% confidence interval 0.0002 to 0.0015) (Fig. S4). There was a moderate positive correlation between the mother *F* and the mean within-mother chick *F* (Pearson r= 0.30, 95% CI 0.05 to 0.52).

### Inbreeding depression on fitness components in adults

In adult males, the return rate was predicted neither by the male *F* nor by the *F* by phenotype interactions, as the interaction models received weak support, and the CIs of these terms overlapped zero (Fig. 2, Tables 2a & 3a). The *F* and tarsus squared model obtained top support (ωAICc=0.28), was 8.8 times more likely than the *F* and tarsus model, and 3.5 times more likely than the null model. The *F* squared models were also well supported (sum of ωAICc= 0.47) but only weakly more probable compared to the null model (Table S2a). The inverse quadratic relationships with the tarsus and *F* indicated that return rates are the highest for males with average tarsi and *F* values, respectively.

**Fig. 1.**
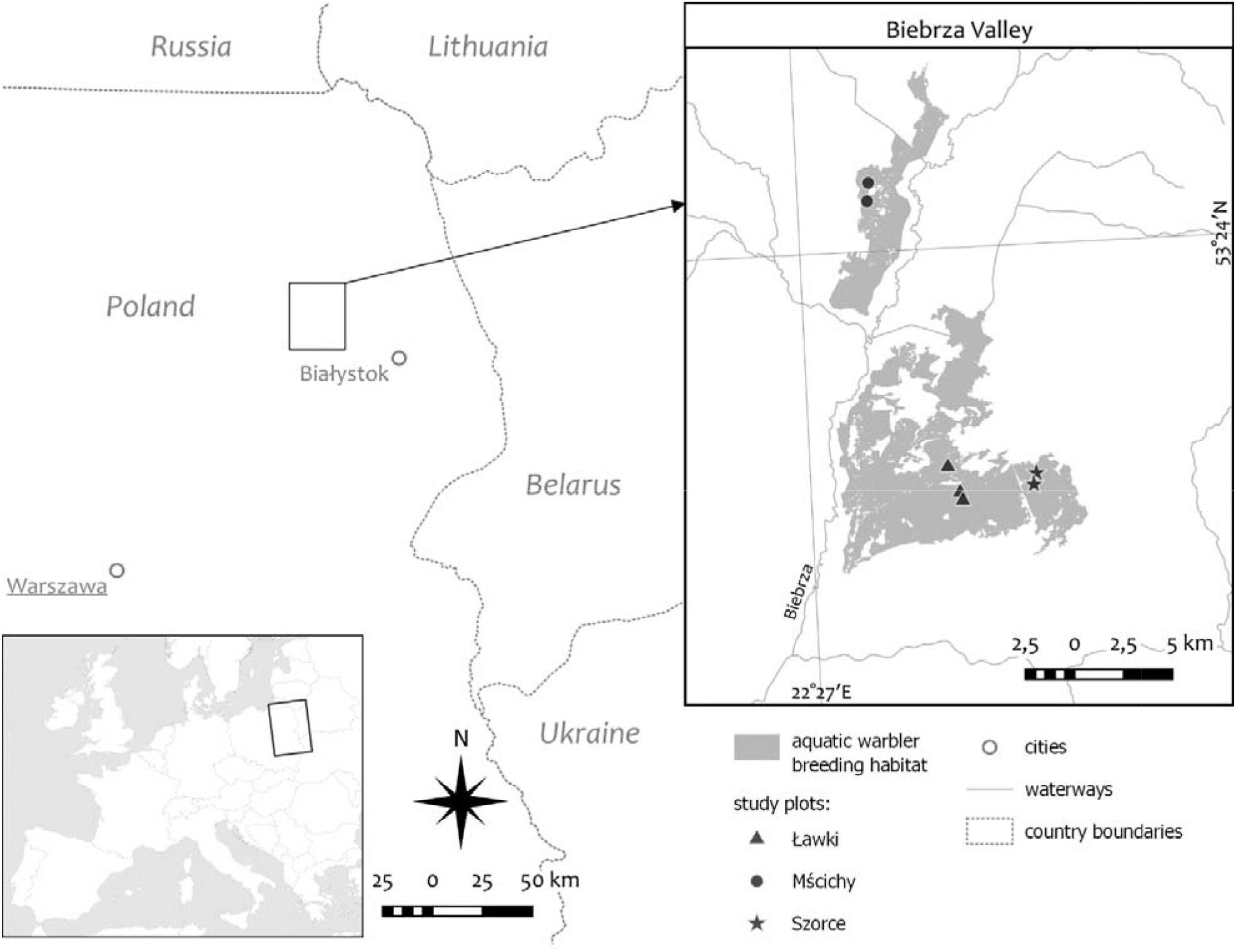
Map of the study area.

**Fig. 2.**
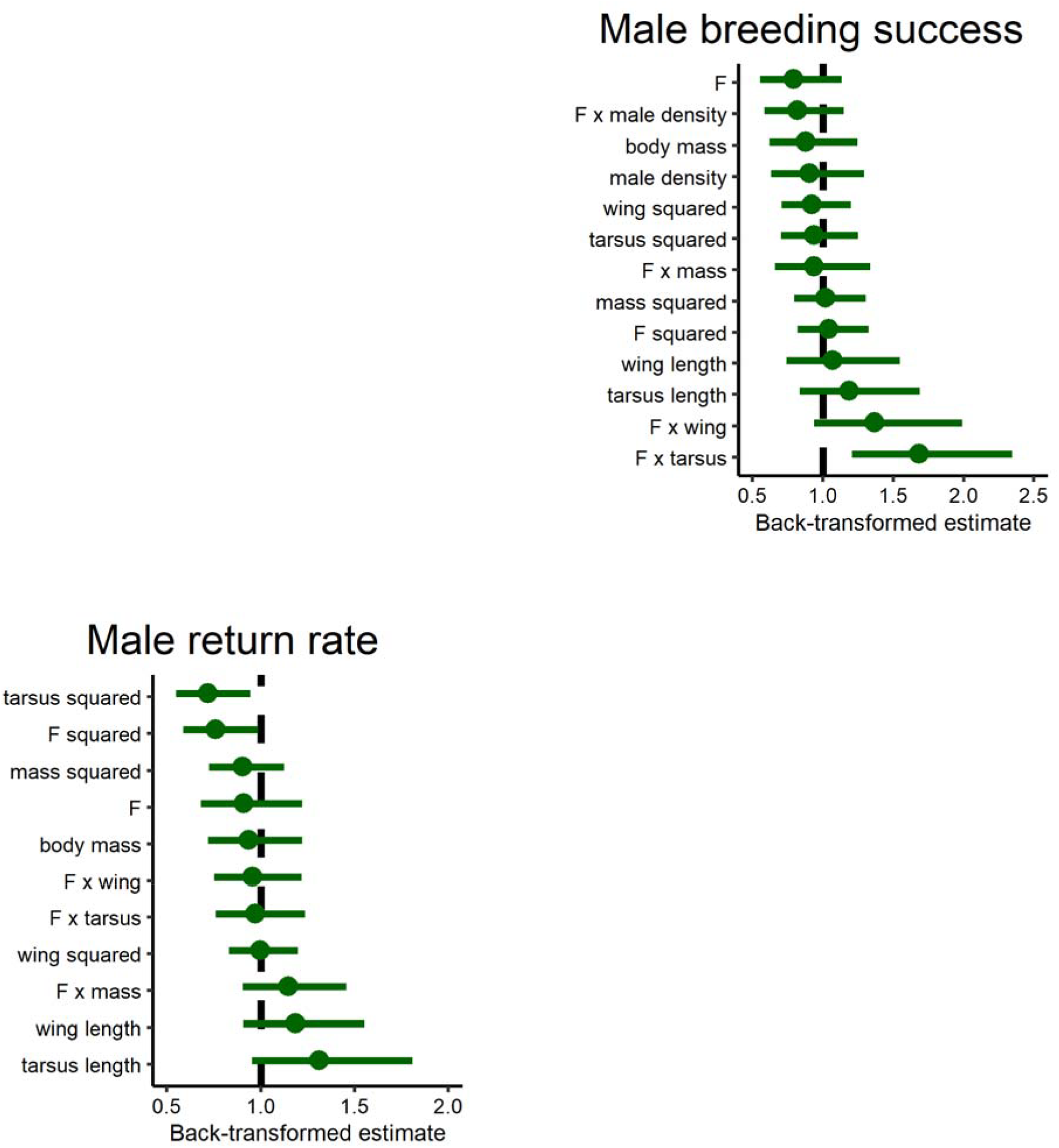
The estimates (denoted by circles) of effects of the male inbreeding coefficient *F*, phenotypic variables and their interaction with *F* on male fitness components: 4-year return rate to breeding grounds (left) and seasonal breeding success (right). The whiskers represent 95% confidence intervals and the vertical dashed line refers to no effect. See Table S3 for the summary of the estimates.

While the male *F* was not a good predictor of the breeding success for average phenotypes and male densities (Fig. 2, Table S3b), the *F* by tarsus interaction model scored the highest (ωAICc=0.47), was 4.9 times more probable than the null model and 16.7 times more probable than the *F* and tarsus model (Table S2b). The remaining interaction effects obtained low support. In males with small tarsi, the *F* estimate was negative, and in males with exceptionally large tarsi it was positive, however, the latter result was based on very few observations and thus uncertain (Fig. 3).

**Fig. 3.**
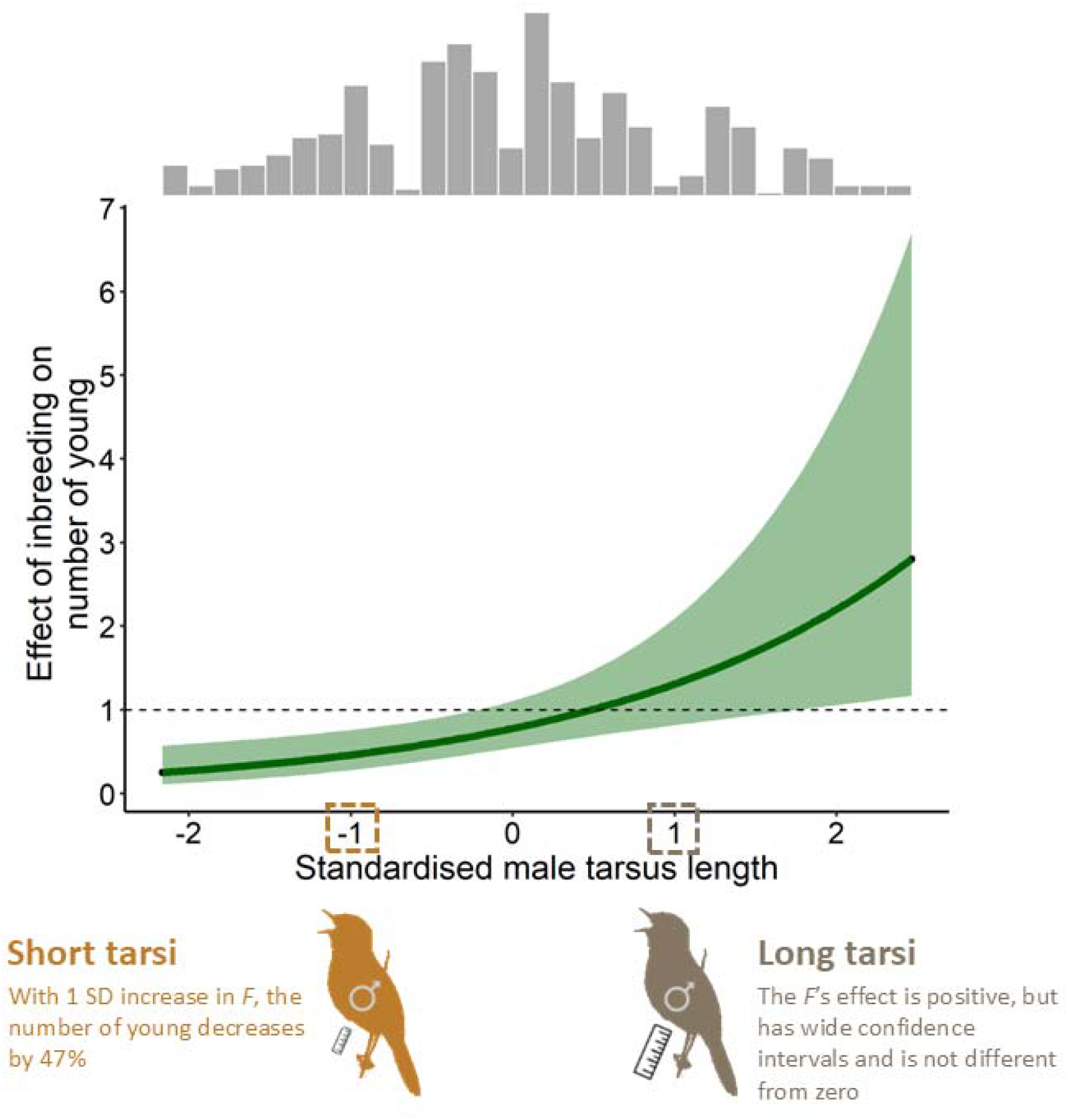
The effect of interaction between the male tarsus length and inbreeding coefficient *F* on the seasonal breeding success. The graph shows how the back-transformed statistical effect of *F* changes against standardised tarsus lengths, and the bands show the 95% confidence interval of the effect. The aquatic warbler pictograms at –1SD and +1SD present examples of how to interpret the graph. The horizontal dashed line corresponds to no effect. The histogram shows frequency of tarsus lengths.

For average covariates, the female *F* relationship with the clutch size was unsupported (Fig. 4, Table S5a). However, the *F* by tarsus interaction model obtained top support (ωAICc=0.52), was 4.3 times more probable than the *F* and tarsus model, and clearly more parsimonious than the null model (Table S4a). The remaining interaction effects were unsupported. In females with small tarsi, the F effect on the clutch size was negative, while in those with the largest tarsi it appeared positive; however, the latter result was based on very few observations and thus uncertain (Fig. 5). The tarsus models ranked the highest (sum of ωAICc 0.99), pointing to the tarsus as the strongest and positive predictor of the clutch size (Fig. 4, Tables S4a&S5a).

**Fig. 4.**
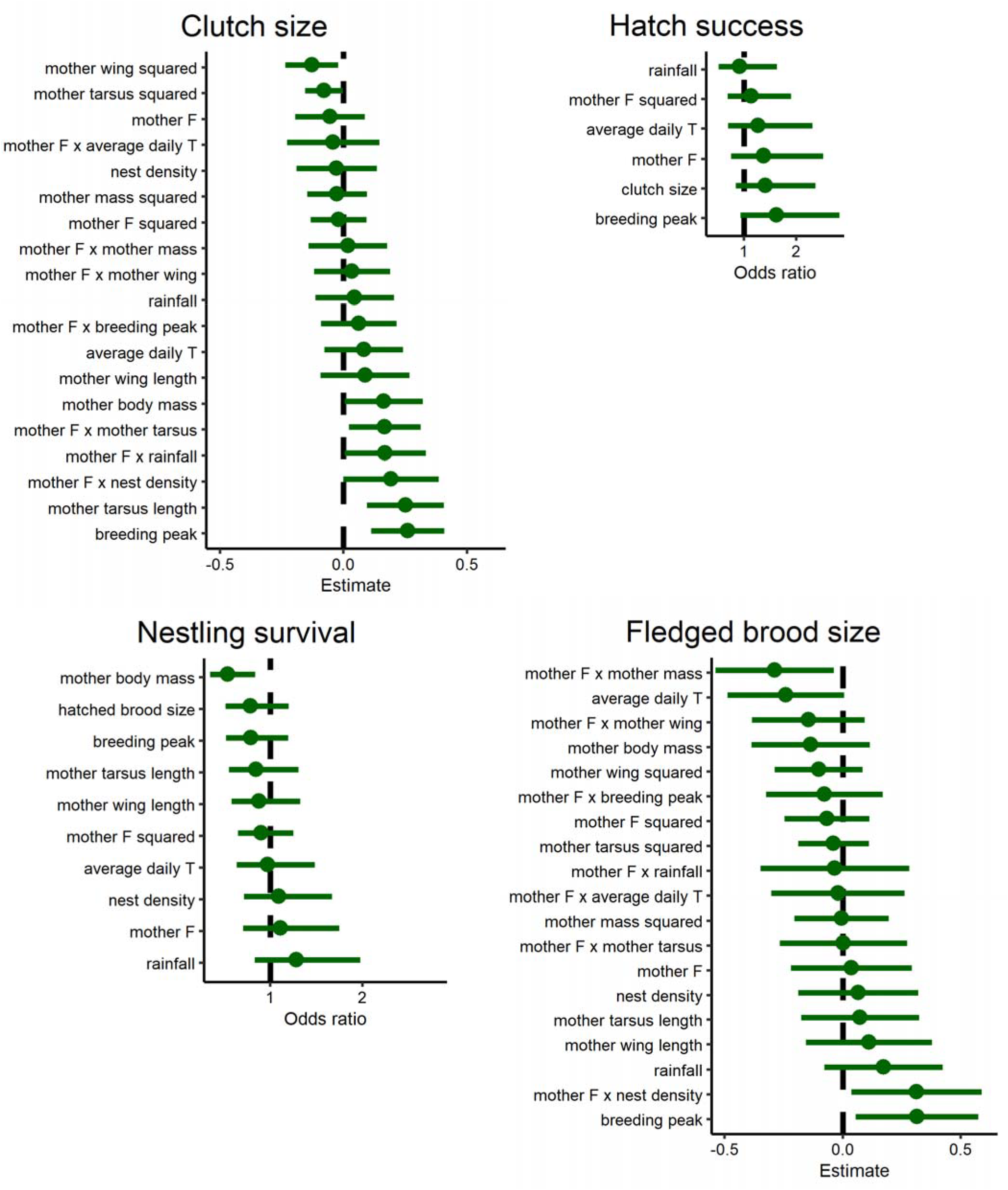
The estimates (denoted by circles) of effects of the mother inbreeding coefficient *F*, phenotypic variables and environmental variables and their interaction with *F* on female fitness components. The whiskers represent 95% confidence intervals and the vertical dashed line refers to no effect. See Table S5 for the summary of the estimates.

**Fig. 5.**
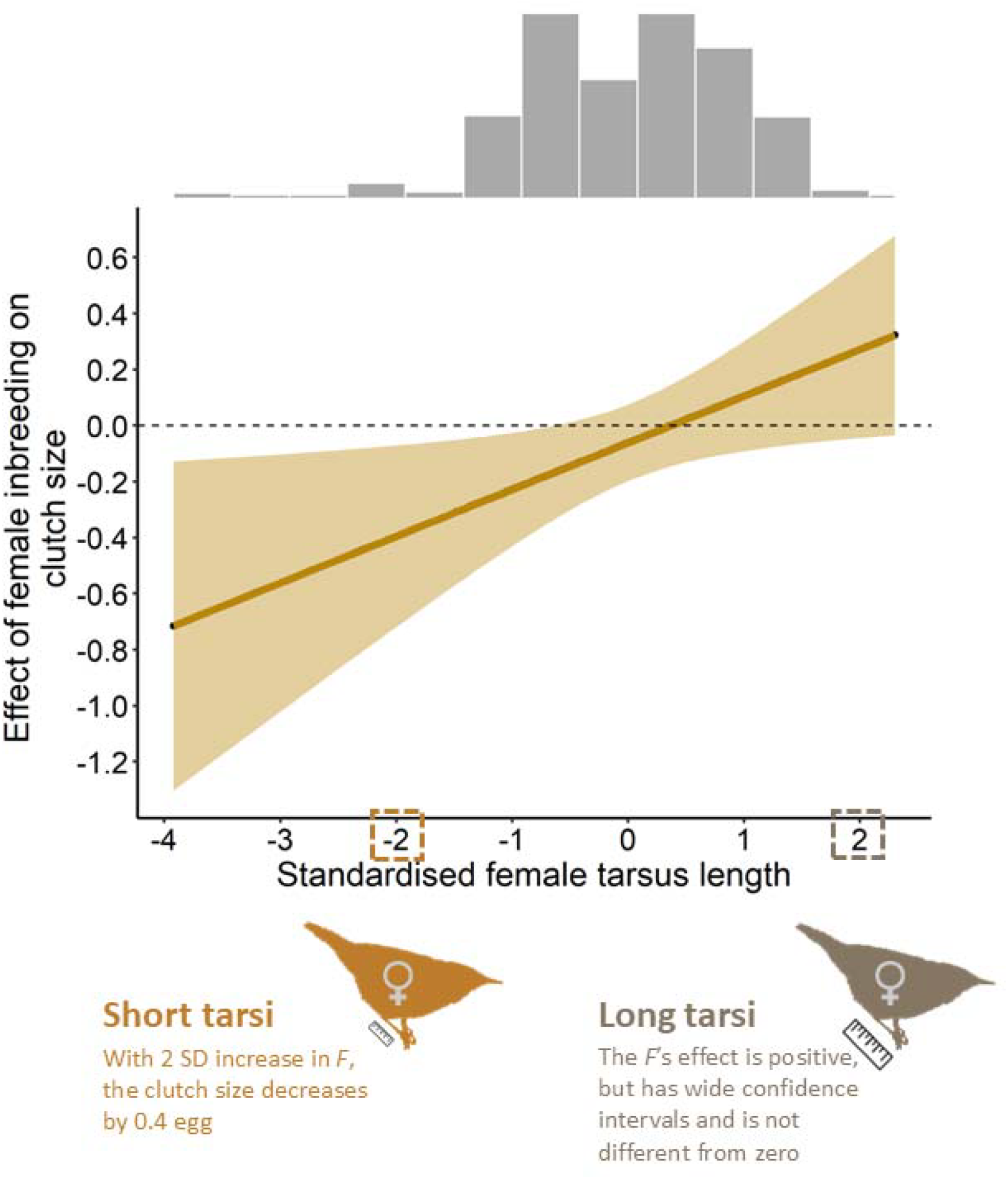
The effect of interaction between the mother tarsus length and inbreeding coefficient *F* on the clutch size. The graph illustrates how the statistical effect of the mother *F* changes along the value of the tarsus. The aquatic warbler pictograms at –2 SD and +2 SD present examples of how to interpret the graph. The bands show the 95% confidence interval of the effect, the horizontal dashed line corresponds to no effect, and the histogram shows the frequency of tarsus lengths.

Mother *F* showed no clear relationship with the hatching success and nestling survival (Fig. 4, Tables S5b-c). For the hatching success, the null model scored the highest (ωAICc=0.21) and support was spread between several models (Table S4b). For the nestling survival, the *F* and body mass model obtained clear support from the data (ωAICc=0.70) (Table S4c), showing that mother body mass was a strong negative correlate of nestling survival (Fig. 4, Table S5c).

Similarly, the relationship between the mother *F* and fledged brood size was not supported for average phenotypic and environmental covariates (Fig. 4, Table S5d). The null model ranked the second and the *F* by body mass interaction model was 4.4 times more probable than the *F* and body mass model, however, it was nearly as parsimonious as the null model, indicating very low support (Table S4d). The remaining interaction effects were not supported. The breeding peak was a relatively strong predictor of the fledged broood size (Fig. 4, Table S5d).

### Inbreeding depression on nestling tarsus

The 95% credible interval of the model-averaged chick *F* effect spanned zero, indicating no support for an association between *F* and tarsus for average covariates (Table S7). Evidence was spread between several animal models (Table S6). The *F* by rainfall and *F* by temperature interaction models scored best (ωDIC=0.12 and 0.11, respectively). The *F* by temperature interaction estimate was negative, suggesting that the *F* effect becomes more negative with increasing temperatures and varies by year. We obtained low evidence for the remaining interactions (Tables S6 & S7).

### Inbreeding load and change in fitness

For average phenotypic traits, the inbreeding load on male return rates was moderate, with nearly half the number of years returned to the breeding grounds in the most inbred males, compared to the least inbred ones, and the inbreeding load on the male breeding success was high (Table 1). However, as the 95% CIs of the male *F* effects spanned zero, these inbreeding load estimates bear high uncertainty. The males with average small tarsi incurred strong inbreeding load, with almost 90% reduction in the number of young for the most inbred males, compared to the least inbred ones. In contrast, in males with average large tarsi the inbreeding load was negligible (Table 1).

**Table 1.** Inbreeding load (*B*) and change in the fitness components due to inbreeding (*δ*).

| Fitness component | Inbreeding determined in: | | Standardised $B$ | Native scale $B$ | Change in trait $-\delta$ |
| --- | --- | --- | --- | --- | --- |
| Male long-term survival | adult male |  | 0.11<sup>1</sup> | 2.45<sup>1</sup> | -48.3%<sup>1</sup> |
| Male seasonal breeding success | adult male |  | 0.21<sup>1</sup> | 4.87<sup>1</sup> | -69.3%<sup>1</sup> |
| Average small males | adult male |  | 0.40<sup>1</sup> | 9.27<sup>1</sup> | -89.4%<sup>1</sup> |
| Average large males | adult male |  | 0.01<sup>1</sup> | 0.21<sup>1</sup> | -5.0%<sup>1</sup> |
| Clutch size | adult female |  | 0.06 | 1.26 | -5.0% |
| Average small females | adult female |  | 0.12 | 2.83 | -11.5% |
| Average large females | adult female |  | -0.01 | -0.28 | +1.1% |
| Hatch failure | adult female |  | -0.01<sup>2</sup> | -0.17<sup>2</sup> | +3.5%<sup>2</sup> |
| Nestling mortality | adult female |  | 0.01<sup>2</sup> | 0.15<sup>2</sup> | -2.9%<sup>2</sup> |
| Fledged brood size in successful nests | adult female |  | -0.03 | -0.66 | +3.4% |

The $B$ is expressed in lethal haploid equivalents and was calculated from weighted-averaged predictions (Keller & Waller 2002, Box 3, Eqn II) of respective candidate model sets (Tables S2, S4 & S6) on the link scale, which were then weighted-averaged between the study areas, years and breeding peaks (if applicable). The native scale $B$ was calculated by back-standardisation. The $\delta$ is reported as the change in fitness of the most inbred relative to the least inbred individuals in the dataset, and was calculated as above, with the final predictions back-transformed to the response scale. For clarity, $-\delta$ is given, i.e. the minus in front of the $\delta$ means a decrease in the trait value, while the plus indicates an increase.
<sup>1</sup> negative binomial distribution with log-link used931 <sup>2</sup> poisson distribution with log-link used

The female inbreeding load on the clutch size was low for the average covariates. However, mothers with average small tarsi bore higher, although moderate inbreeding load, with about 10% reduction in the clutch size for the most inbred relative to the least inbred females, while in mothers with average large tarsi the inbreeding load was very low (Table 1). The mother inbreeding load on the hatching success, nestling survival and fledged brood size was also low. The overall decrease in fitness between laying a clutch and brood fledging totalled 0.6 haploid lethal equivalents (Table 1).

## Discussion

### Weak inbreeding depression for average phenotypic and environmental variables

For the mean value of the phenotypic traits and male density in the case of the breeding success, inbreeding depression on the long-term return rates and seasonal breeding success of adult males obtained low support from the data. Male aquatic warblers show high post-breeding annual return rates (Dyrcz & Zdunek 1993, Bellebaum 2018), which are comparable to survival rates of other migratory passerines in the central European latitude (Scholer *et al*. 2020). Hence, the 4-year return rate is a strong predictor for the lifespan survival. Given the median return rate of 1 year among the surviving males, most males reproduce over 1 breeding season, indicating that the seasonal breeding success is a proxy for the lifetime reproductive success. Therefore, our results do not suggest that the inbreeding rate in adult males is associated with their adult lifespan nor the lifetime reproductive success.

Avian studies on the inbreeding depression on adult survival have been scarce and showed weak, e.g. *B*= 1.7 (Keller 1998), moderate, e.g. *δ*= 49% (Harrisson *et al*. 2019) or strong effects of inbreeding, e.g. up to *B*= 7 in the house sparrow (*Passer domesticus*) (Niskanen *et al*. 2020). Evidence for male inbreeding effects on the annual reproductive success has also been variable, ranging from weak support (Sin *et al*. 2021), through moderate effects, e.g. *δ*= 42% (Harrisson *et al*. 2019) to clearly negative inbreeding depression in the house sparrow, *B*= 6 (Niskanen *et al*. 2020). The *B*= 4.9 and *δ*= 69% that we obtained for aquatic warbler males appear to be high inbreeding load on the annual reproductive success, however, these estimates bear high uncertainty.

Similarly to males, for the mean value of the phenotypic and environmental covariates, we found weak support for the relationship between an adult female’s inbreeding rate and her short-term fitness components. These results do not support mother inbreeding depression on offspring production from clutch laying until fledging in the aquatic warbler. However, it is the earliest developmental stages that could respond more negatively to inbreeding (Noordwijk & Scharloo 1981, Hemmings *et al*. 2012b), and it is inbreeding of the offspring that could affect its survival more strongly.

Previous studies on birds found the parental inbreeding to be little related to egg production (Keller 1998, Szulkin *et al*. 2007, Harrisson *et al*. 2019). In the takahe (*Porphyrio hochstetteri*), parental inbreeding was unassociated with hatching rates (*B*= –0.7) and fledging success (*B*= 3.3) (Grueber *et al*. 2010), but in another study the latter decreased by 30% in mothers with the pedigree *f*>0 (Jamieson *et al*. 2003). Negative effects of maternal inbreeding were demonstrated in two studies on passerine birds for the hatching success (*δ*= 20-36%) and fledgling production (*δ*= 26%) (Keller 1998, Harrisson *et al*. 2019). In our study, although the clutch size manifested the highest inbreeding load of the female fitness components studied, it was relatively low, and so was the inbreeding load from clutch laying to brood fledging.

Neither did we find unambiguous evidence that, for average values of the phenotypic and environmental covariates, and while accounting for mother inbreeding, the offspring inbreeding rate is associated with the tarsus length, and this association remained constant across the three nestling ages. This does not support inbreeding depression on the skeletal body size in aquatic warbler nestlings, for typical individuals and conditions. In contrast, in the collared flycatcher (*Ficedula albicollis*), inbred fledglings had smaller tarsi compared to outbred ones (Kruuk *et al*. 2002), but no such relationship was found in another species (Sin *et al*. 2021).

### Effects of inbreeding depend on body size in adults and on weather in offspring

We demonstrated that inbreeding depression on the male seasonal breeding success and clutch size is conditional on the tarsus length of males and females, respectively. In birds, tarsus length is a proxy for the structural body size (Rising & Somers 1989, Senar & Pascual 1997), implying that the inbreeding costs on these fitness-related traits increase in small-bodied individuals, relative to large-bodied individuals. In some passerines, the tarsus or a body size index is positively associated with the clutch size in females (Alatalo & Lundberg 1986, Sedinger *et al*. 1995, Garamszegi *et al*. 2004, this study) and with sperm quality and functionality traits and testis size in males (Brown & Brown 2003, Forstmeier *et al*. 2017), which translates into sperm production (Parker & Pizzari 2010, Hayward & Gillooly 2011). Therefore, it could be more difficult for small inbred individuals to compensate for their lower fecundity traits, than it is for large inbred ones.

The inbreeding depression on the breeding success in the small-bodied males appears strong. For example, the *B* of 9 and ~89% decrease in the male annual reproductive success are comparable to the high inbreeding depression in the house sparrow (Niskanen *et al*. 2020). In small-bodied females, the inbreeding load on clutch sizes (*B*= 2.8) and the decrease in the clutch size produced (~11%) were moderate and did not translate to decreased fledgling production. Our finding of inbreeding depression on the clutch size in small females stands in contrast with previous studies (Keller 1998, Szulkin *et al*. 2007, Harrisson *et al*. 2019).

We did not obtain firm support for environmental dependence of the chick inbreeding effects on tarsus, except for a weak interaction with the average daily temperature. This interaction suggests that chick *F* was more negatively associated with the tarsus in higher temperatures in the preceding days. As temperature was standardised within year, breeding peak and day, its effect was independent of these factors. Elevated temperatures could correlate with lower abundance or availability of prey, and thus more inbred chicks could cope worse with being fed less during warmer days. As the effect of nestling *F* varied between the two study years, the magnitude and direction of inbreeding effects on nestling body size could differ between breeding seasons.

### Conclusions and conservation implications

We sought to determine the magnitude of the inbreeding depression on fitness components and nestling growth in a threatened passerine, the aquatic warbler, and to inspect whether the inbreeding effects on these traits are modulated by the phenotype and environment. We demonstrated, to our knowledge for the first time, that in birds, inbreeding depression on adult fitness components could depend on the body size, with smaller individuals paying higher costs of inbreeding. This suggests that birds raised in poor environmental conditions, which negatively affect the adult body size (De Kogel 1997, Searcy *et al*. 2004, Cleasby *et al*. 2011) could have lower fitness if they are inbred. Our study also points out that accounting for the body size (and possibly other phenotypic and environmental variables) allows to determine inbreeding depression more precisely. This is of considerable importance especially in the case of species of conservation concern, where nuanced knowledge on inbreeding depression will be informative for adequate estimation of extinction risk. Finally, our observation that nestling inbreeding association with tarsus is conditioned by the temperature and year partly corroborates that unfavourable environmental conditions could exacerbate inbreeding depression. It also implies that survival to reproductive maturity, of which the chick tarsus length is a proxy, could be negatively affected for nestlings growing in excessively warm ambient temperatures.

For the aquatic warbler, our study shows that for average phenotypic and environmental covariates, inbreeding effects on the studied fitness traits are weakly supported. However, we were able to estimate inbreeding depression on fitness traits only in adults. Weakly negative inbreeding effects on short-term fitness components can accumulate to high inbreeding load over the complete life history continuum (Szulkin *et al*. 2007, Grueber *et al*. 2010, Harrisson *et al*. 2019, Niskanen *et al*. 2020). Future studies on the inbreeding depression in the aquatic warbler should include the missing fitness components, e.g. survival from viable egg to sexual maturity, juvenile survival and adult female survival. Notwithstanding, our results will be informative for a population viability analysis accounting for inbreeding depression (O’Grady *et al*. 2006, Nonaka *et al*. 2019) and for conservation actions such as translocation.

## Acknowledgements

Grzegorz Kiljan, Pedro Costa, Piotr Guzik, Marta Celej, Julien Foucher, Beata Głębocka, Aneta Gołębiewska, Maciej Kamiński, Agnieszka Kuczyńska, Romuald Mikusek, Łukasz Mucha, Felix Närmann, Edyta Podmokła, Aneta Rybińska, Paulina Siejka, Michał Walesiak and Laura Zani assisted in the field. Gugny Field Station of the Białystok University provided accommodation and workspace during fieldwork. Susan Mbedi, Ismael Reyes and Katarzyna Dudek supported the preparation and sequencing of RAD libraries, and Wiesław Babik provided lab space and equipment. The labwork and bioinformatic analysis were partially funded by the German Federal Ministry of Education and Research (BMBF, Förderkennzeichen 033W034A). The sequencing was performed in cooperation with the Competence Center for Genome Analysis at the Christian-Albrechts-University in Kiel and at the Berlin Center for Genomics in Biodiversity Research. The study was funded by the Polish National Science Centre’s grant no. 2016/20/S/NZ8/00434 awarded to Justyna Kubacka.

