## SupplementaryMaterial1 for "It is hard to be small: Inbreeding depression depends on the body size in a threatened songbird"

**Supplementary Online Resource 1**

**Detailed description of RAD-seq population libraries**

The nine population libraries were processed in the following batches and libraries: Batch 1) 88 females (Cat1 – 30 individuals, Cat2 – 29 individuals and Cat3 – 29 individuals); Batch 2) 243 offspring and 45 males with song recordings (Lib1 and Lib2 – 144 individuals each); Batch 3) 11 females (Lib3); and Batch 4) 82 offspring (including two samples that failed in Lib1 & Lib2) and 131 males (Lib4, Lib5 and Lib6 – 71 individuals each). All the libraries were balanced with respect to the three sampling locations.

The population libraries were prepared according to the reference libraries protocol, with the exception that: 1) a size range of 350-500 bp was excised with Blue Pippin for libraries Lib1-Lib6; 2) the size selection for Lib4, Lib5 and Lib6 was done after the indexing PCR and lacked the step of a single-cycle PCR adding an Itru5-8N primer; 3) for Lib4, Lib5 and Lib6 purification was carried out with AmPure beads; and 4) the number of cycles in the final indexing PCR was 12.

Sequencing was done as follows: libraries Cat1, Cat2, Cat3 and Lib3 – lllumina NextSeq MidOutput 300 cycles; Lib1 and Lib2, as well as Libs 4-6 – Illumina NovaSeq 6000 300 cycles, one lane of the SP flow cell.

**Table S1**. Number of sequenced reads obtained by library (a) and number of loci and SNPs remaining after each filtering step of the combined sequences (b).

a)

| Lib1 | Lib2 | Cat1 | Cat2 | Cat3 | Lib3 | Lib4 | Lib5 | Lib6 |
| --- | --- | --- | --- | --- | --- | --- | --- | --- |
| 218.8 M | 199.7 M | 22.9 M | 23.4 M | 23.5 M | 10.8 M | 157 M | 135 M | 98 M |

b)

| STACKS:  gstacks | STACKS:  populations   - r=0.8 - p=1 - min-maf=0.005 - write 1st SNP only | Vcftools:   - minDP 10 - max-missing 0.8 | Vcftools:   - max-meanDP 115 | Vcftools:   - hwe 0.05 - FDR correction | Vcftools:   - remove one locus from each pair with r^2^>0.3 | Vcftools:   - maf 0.3 - max-missing 0.9   (for parentage analysis) |
| --- | --- | --- | --- | --- | --- | --- |
| 6267 RAD-loci | 4179 SNPs | 3892 SNPs | 3886 SNPs | 3514 SNPs | 2948 SNPs | 333 SNPs |

**Table S2**. A summary of the fitness-components measured in adult male and female aquatic warblers.

| fitness component | *N* | mean ±SD | median (range) | % |
| --- | --- | --- | --- | --- |
| adult male return rate over 4 years  ringing year 2017  ringing year 2018  ringing year 2019 | 166^1^  77^1^  68^1^  21^1^ | 0.5 ±0.8  0.5 ±0.9  0.5 ±0.9  0.3 ±0.5 | 0 (0-4)  0 (0-4)  0 (0-4)  0 (0-1) | N/A |
| adult male seasonal breeding success  2017  2018 (including 24 males ringed in 2017) | 169^1^  77^1^  92^1^ | 1.0 ±2.2  1.1 ±2.2  1.0 ±2.2 | 0 (0-12)  0 (0-11)  0 (0-12) | N/A |
| clutch size  2017  2018  2019 | 77^2^  33^2^  24^2^  20^2^ | 4.8 ±0.7  4.8 ±0.6  4.8 ±0.7  4.5 ±0.9 | 5 (2-6)  5 (3-6)  5 (4-6)  5 (2-6) | N/A |
| hatch success  2017  2018  2019 | 358^3^  160^3^  116^3^  82^3^ | N/A | N/A | 94  91  97  95 |
| nestling survival  2017  2018  2019 | 285^4^  129^4^  101^4^  55^4^ | N/A | N/A | 83  80  86  85 |
| fledged brood size in successful nests  2017  2018  2019 | 95^5^  49^5^  32^5^  14^5^ | 3.8 ±1.2  3.7 ±1.2  4.1 ±1.1  3.6 ±1.5 | 4 (1-6)  4 (1-6)  4 (1-6)  4 (1-6) | N/A |

*N* – sample size, SD – standard deviation, N/A – not applicable

^1^excluding records missing tarsus, mass or *F* (*N*= 12)

^2^excluding records missing tarsus, mass, wing or *F* (*N*= 7)

^3^excluding records missing mother *F* or hatch success (*N*= 41)

^4^excluding records missing nestling survival or prey abundance (*N*= 54)

^5^excluding records missing mother *F*, tarsus, wing, mass or prey abundance (*N*= 7)

**Fig. S1.** Histogram of loci-pairwise r^2^ values (a) below 0.1 and (b) above 0.1.

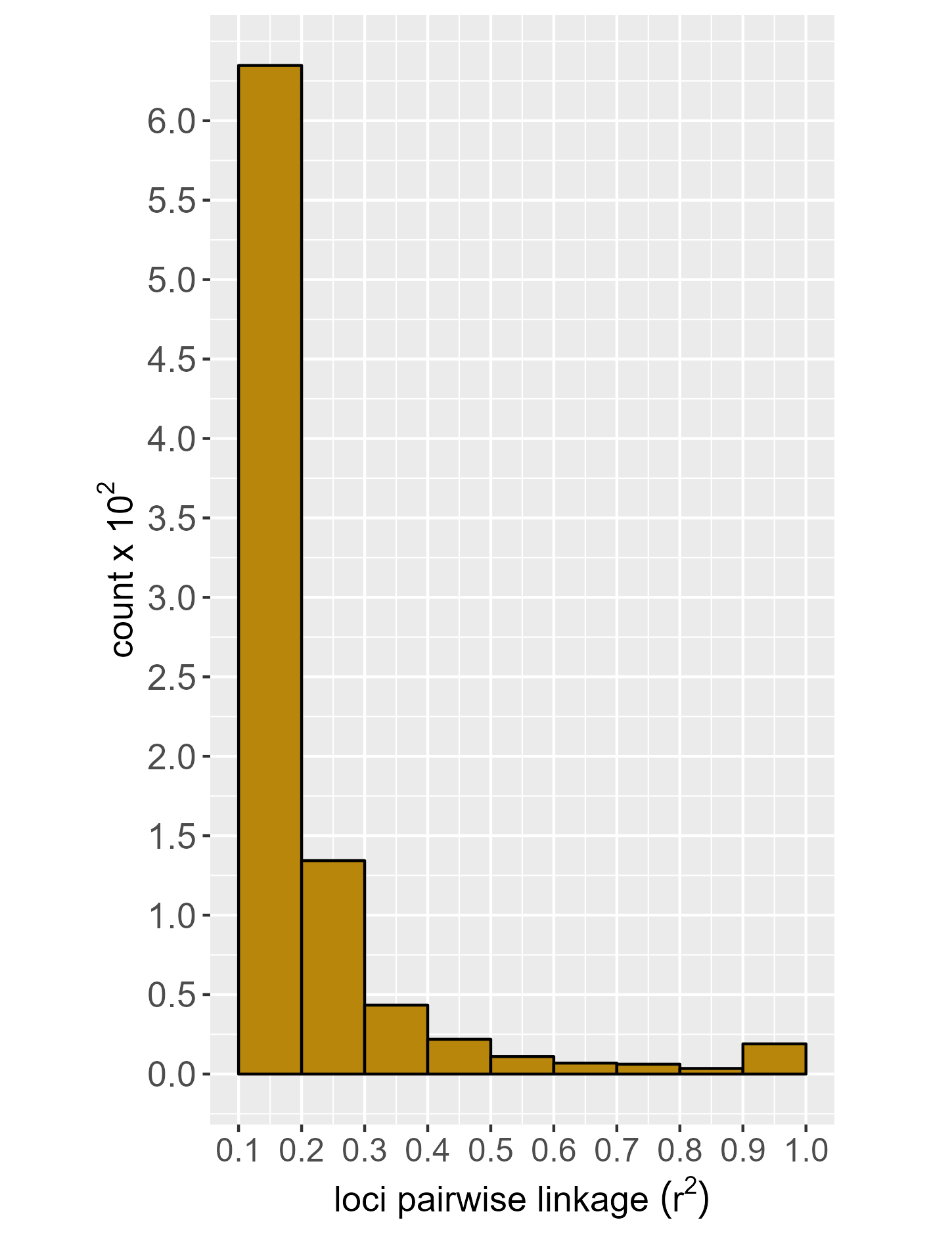

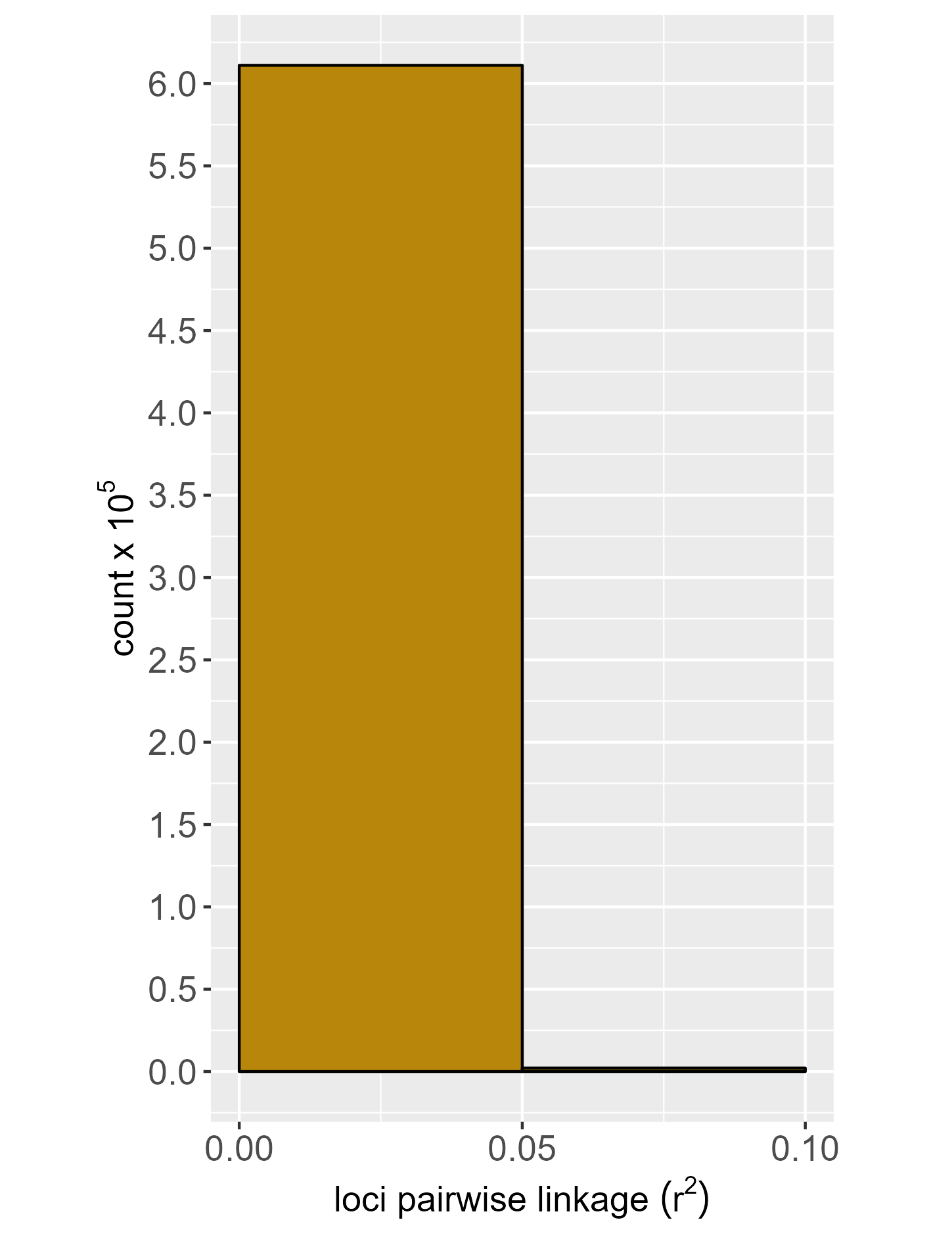
**(a)	(b)**

**Fig. S2.** Histogram of (a) locus depth, (b) individual depth and (c) individual missingness, after removal of loci not passing the minimum genotype depth, locus missingness, maximum mean locus depth, HWE equilibrium and linkage disequilibrium filters. The green vertical dashed line denotes the mean. The mean ±SD read depth per locus and individual was 77.5 ±17.0 and 77.5 ±38.8, respectively, and the median ±SD missingness per individual was 0.014, range 0.001 to 0.725.

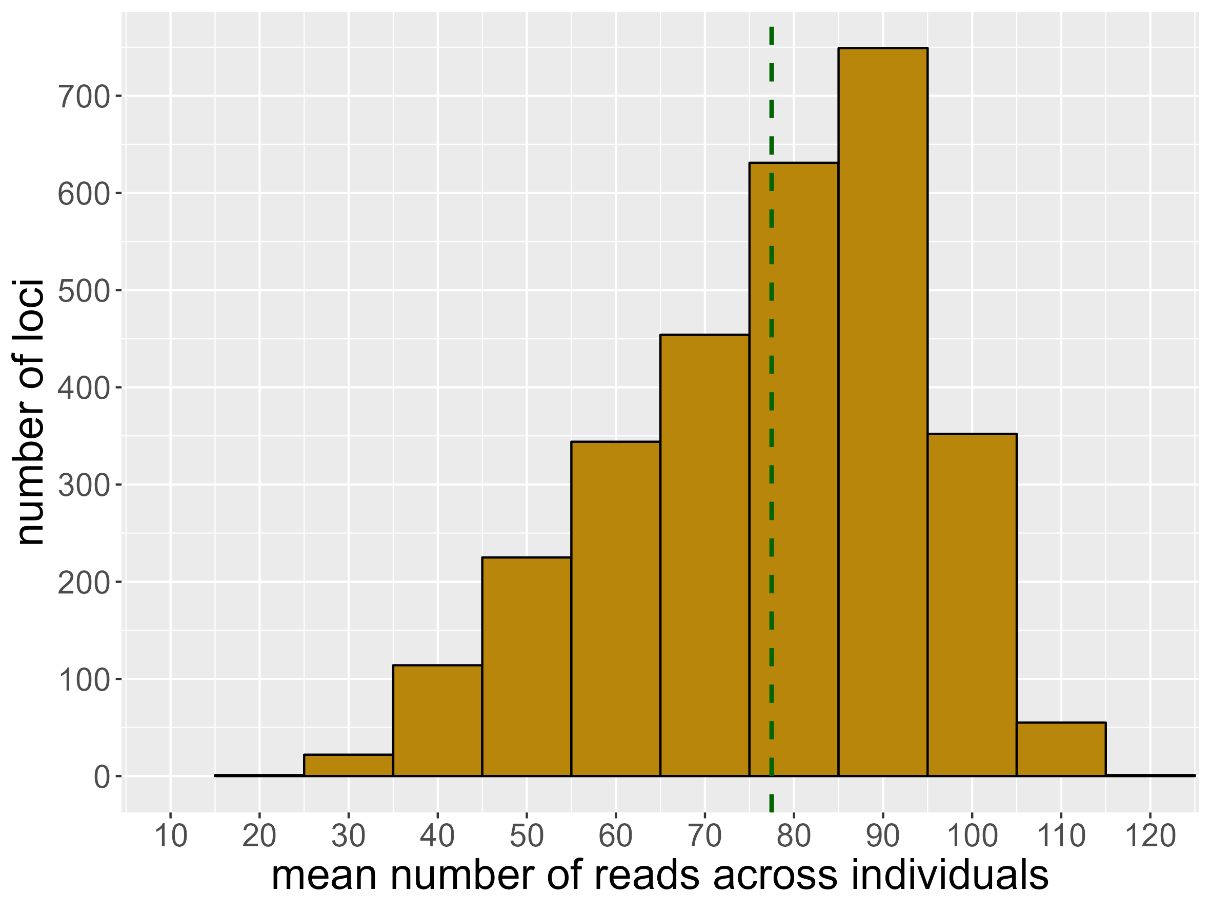

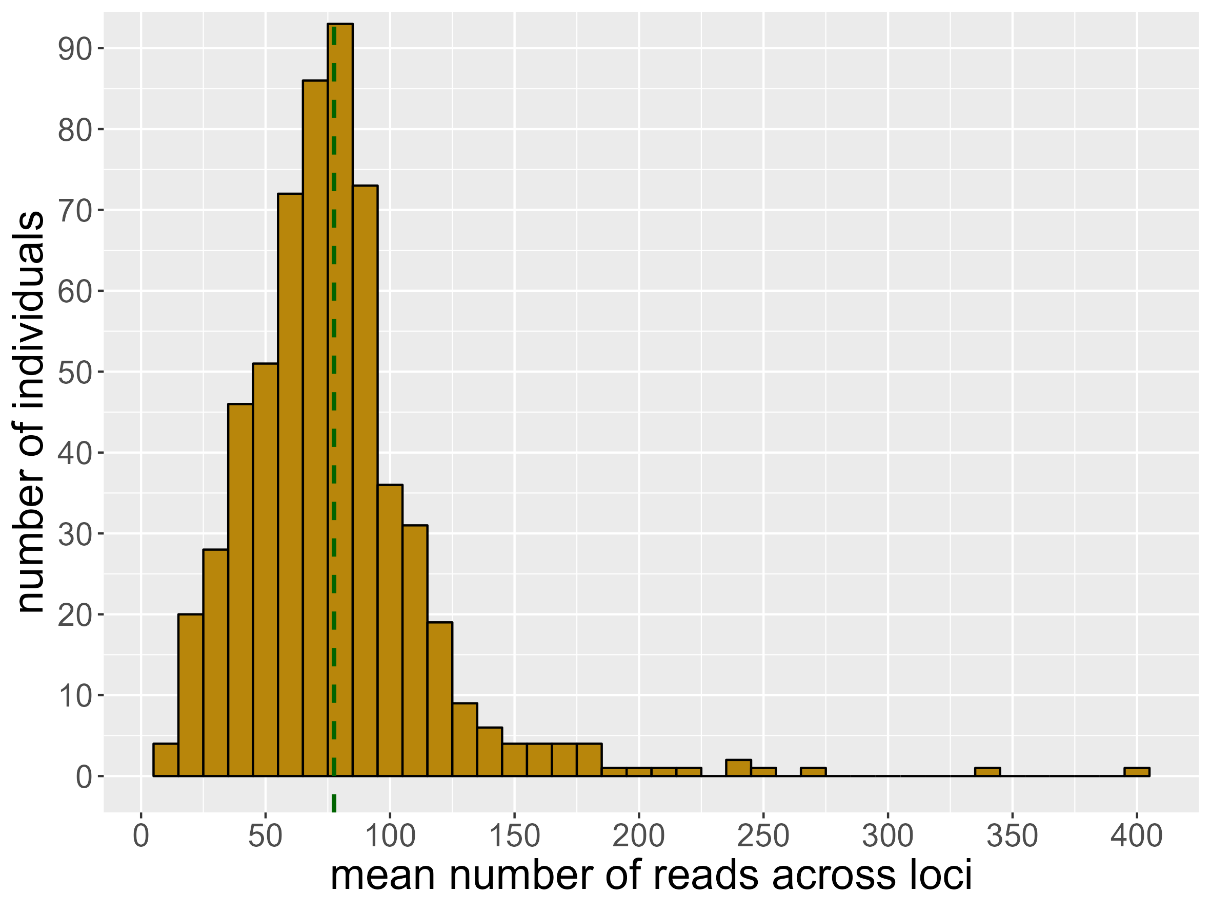
(a)	(b)

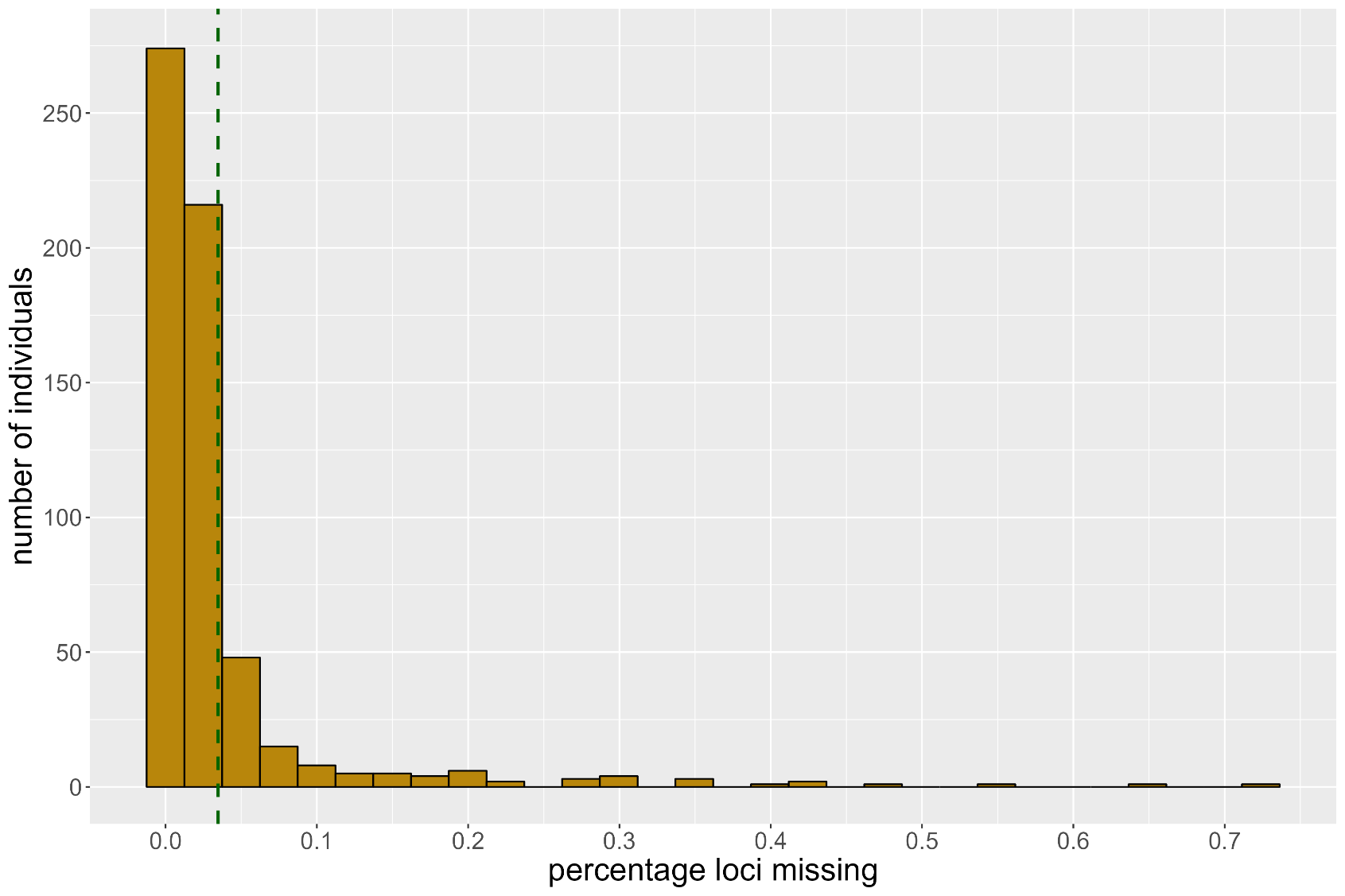
c)

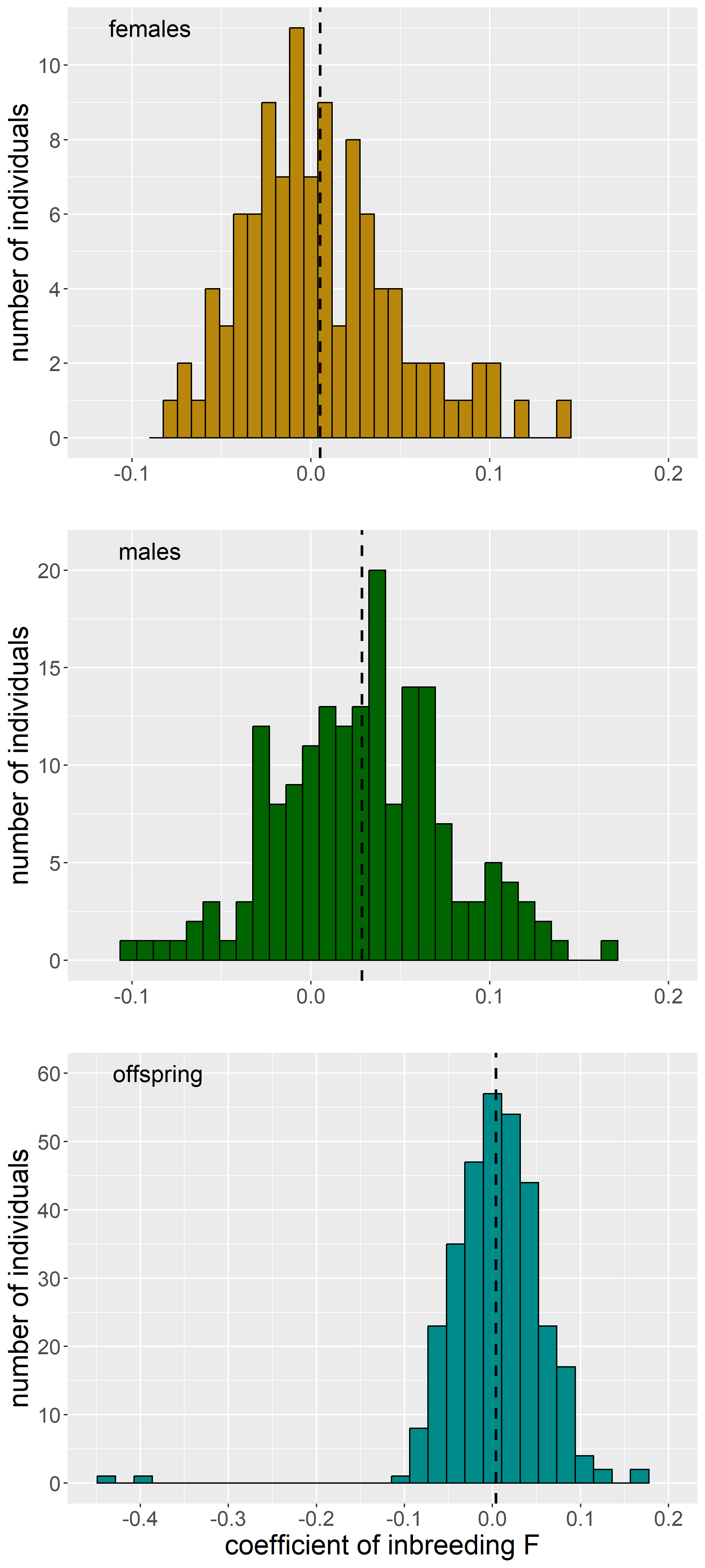
**Fig. S3.** Histograms of inbreeding coefficients (*F*). The vertical dashed lines denote the mean.

**Fig. S4.** Histogram of the identity disequilibrium parameter *g_2_*. The dashed line represents the mean and whiskers denote the 95% confidence interval.

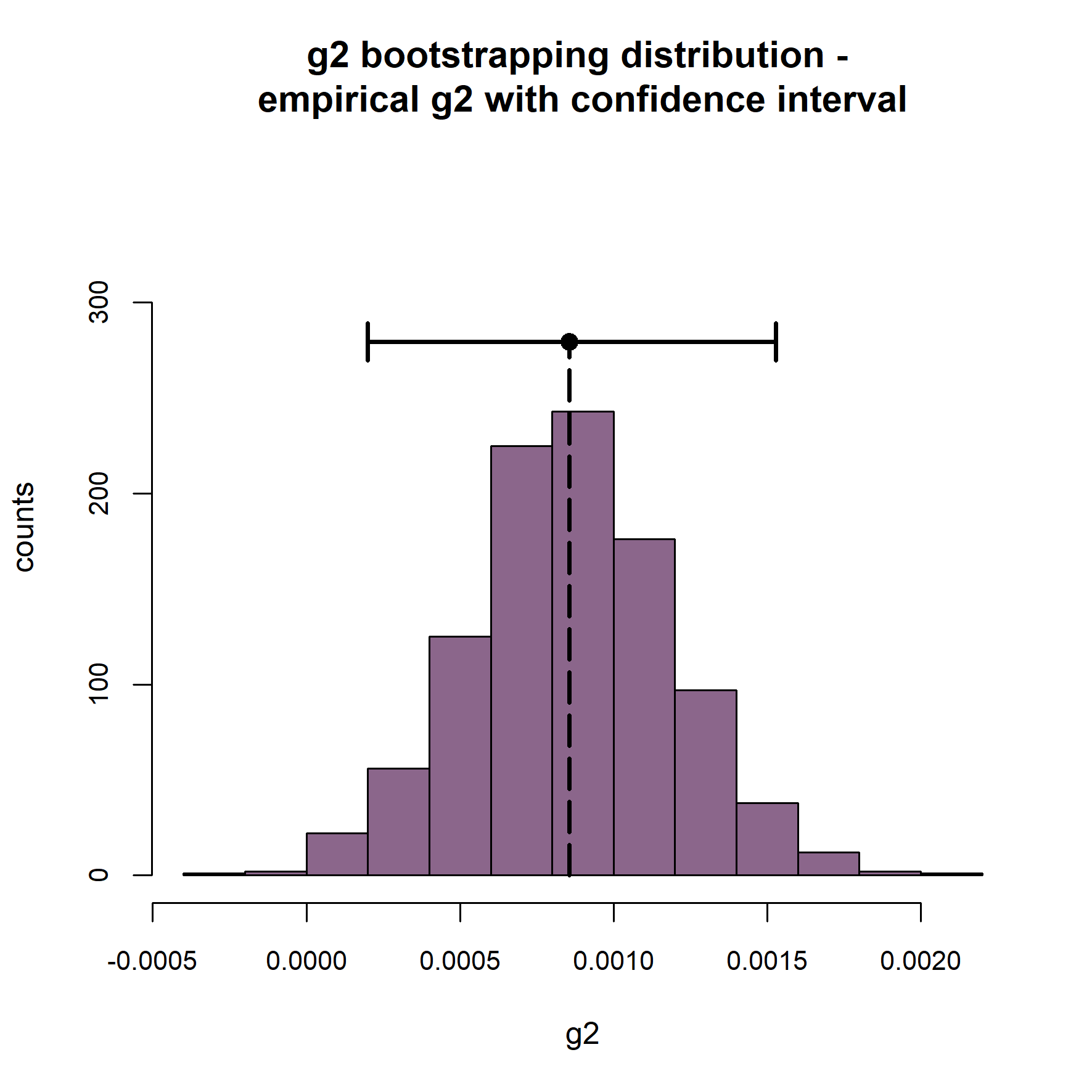
